# Benchmarking siRNA Prediction: The Role of Representation and Validation Strategies

**DOI:** 10.64898/2026.05.12.724560

**Authors:** Aparajita Karmakar, Abdulhamid Merii, Angus Weir, Grzegorz Kudla, Mark Basham, Alex Lubbock

## Abstract

Small interfering RNAs (siRNAs) offer transformative potential for targeted therapeutics, yet the design of highly effective and non-toxic candidates is hindered by the risk of off-target effects and RNA instability. A critical flaw in in silico prediction models is pervasive data leakage in cross-validation protocols, which artificially inflates performance metrics and produces untrustworthy results. To address this, we developed a rigorous framework that eliminates data leakage through strict cross-validation, leverages z-curves (3D representations of RNA physico-chemical properties) for context-aware sequence encoding, and identifies key sequence regions critical for efficacy. Our model achieves an AUC of 0.845 on leakage-free validation, surpassing prior work at 380x faster computation speed, demonstrating that superior representation trumps model complexity. Crucially, we demonstrate how experimental variability and cross-validation choices directly impact model reliability, establishing the first benchmarked methods for robust siRNA efficacy prediction. This work provides a foundation for trustworthy sequence design and validation in RNA therapeutics.

## Introduction

RNA interference (RNAi) is expected to become an essential approach in treating several diseases such as, haemato-oncological, cardiovascular, and neurodegenerative conditions. Its mechanism of action is based on post-transcriptional gene silencing [1]. Small interfering RNAs (siRNAs) are 19-23 nucleotide long double-stranded RNA macromolecules which function in tandem with the RISC (RNA-induced silencing complex), leading to the degradation of the passenger strand (the messenger RNA target). The guide strand guides the RISC complex to the fully complementary target region on the mRNA to which the siRNA binds and cleaves. This natural mechanism can be exploited for therapeutic benefit by introducing synthetic siRNAs into target cells. Therapeutic approaches utilizing siRNA’s silencing mechanism offer numerous benefits compared to protein therapies and small molecule approaches. siRNA targets otherwise “undruggable” locations by directly degrading the target mRNA irrespective of the gene products and offers high selectivity for precise targeting of specific mRNA molecules in disease processes. The synthesis of siRNA molecules is straightforward, allowing for quick modifications and production for therapeutic uses [1].

However, significant challenges remain in developing viable siRNA-based therapeutics. The difficulty of siRNA design is highlighted by the limited number (five) of approved siRNA-based therapeutics for clinical use to date [2]. Despite the potential therapeutic advantages of siRNAs, numerous challenges such as siRNA instability, off-target effects, and immune reactions have significantly hindered the utilisation of siRNA-based treatments [3]. Several factors play an important role in the knockdown effect, including binding affinity (structural motifs), chemical modifications, multiple gene transcripts, and different concentration levels [3], [4]. Predicting the efficacy of siRNA candidates using in silico methods can allow for the selection of siRNAs with high efficacy and low off-target effects. This predictive approach streamlines the subsequent in vitro production of siRNA molecules, optimising their potential for therapeutic use and advancing the development of RNAi-based therapies.

Several methods have been proposed in the literature previously for knockdown efficacy prediction, including traditional machine learning methods as well as advanced methods such as convolutional neural networks [5], transformers [6], [7] and graph neural networks [8], [9]. Earlier experiments tested several classical ML methods like Support Vector Machines and ensemble methods such as Random Forest which often demonstrated better performances than Artificial Neural Network(ANN) models [10]. One of the earliest implementations of ANN was called Biopredsi by Huesken et al. [11]. This work served as a benchmark for future works. While there were more explorations around ANNs, Convolutional Neural Networks (CNN) and Graph Neural Networks (GNN) emerged to be better alternatives. In Han et al’s work, the authors have proposed a CNN model that achieved high accuracy by capturing non-linear relationships within data [5]. On the other hand, La Rosa et al has implemented Graph Neural Network on sequence and thermodynamic features to capture hidden relationships between siRNA and mRNA [8]. Finally, in 2024, Liu et al implemented Transformer Encoder Layers, an attention-based approach to predict siRNA inhibition and off-target effects [6].

However, one of the major drawbacks of the previous works were the cross-validation methods used. There is no standardised cross-validation method adapted in the literature. One of the most used methods is the K-fold cross-validation which divides the data into *K* equal sized parts and in each iteration trains on the *K-1* parts and tests on the remaining one part. The problem with this validation technique is the repetition of targets/genes within the folds that can lead to potential bias in the evaluation process leading to unreliable metrics. This issue was also recently highlighted by Long et al in the siRNADiscovery paper [9]. In order to overcome the bias, they have suggested splitting the data by target/siRNA groups. This ensures a more rational and unbiased evaluation method, however, still does not solve for bias in the choice of candidates for train/test.

Finally, it is important to identify features that contribute towards knockdown prediction. These features can be extracted manually or through the feature extraction layers of deep learning models. Previous works have mostly often relied on one-hot or k-mer representations and thermodynamic features [8], [6]. La Rosa for instance, have used k-mers to represent siRNAs and mRNAs. However, not only are the k-mer vector sizes variable, but they have also used the entire mRNA instead of the binding region, which can lead to increase in dimensions as well as redundancy. Moreover, the k-mer based method fails to capture other chemical features of the sequence very well. Some of them have also considered other features such as G/C content and binding rules [9]. siRNADiscovery has identified key features such as G/C content, binding probabilities and interactions with the Ago2 protein (which facilitates mRNA cleavage) among others. These features have been manually extracted to create the feature space. Some of the more recent works have used RNA foundation model embeddings as well [6], [7].

RNA-FM, trained on 23 million non-coding RNA (ncRNA) sequences, effectively captures structural and functional features from sequence data; however, it is primarily optimized for non-coding RNAs. Sequence representation is one of the major factors that could impact the model’s performance, making it important to identify the best embedding or set of features to represent the sequences.

In this study, we explore multiple sequence representation strategies combined with classical machine learning models to develop a fast and accurate predictor of siRNA activity. To ensure an unbiased evaluation, we aim to benchmark a stricter cross-validation scheme. Following model development, we perform feature analysis to identify the key determinants driving the knockdown process.

## Materials and methods

### Datasets

We have collected our datasets from La Rosa’s work [8]. This dataset has 2816 siRNA-mRNA interactions from [11], [12], [13], [14], [15] and [16]. An optimal efficacy prediction model should be capable of learning from and making predictions across various experiments. However, it is important to note that the six experiments above have different protocols, while the siRNAs have different modifications and concentrations. This is why we will present most of our results using a subset of the above datasets, the Huesken’s dataset, as it represents over 85% of the entire dataset. By using Huesken’s subset, we shall look at our model’s performance across homogeneous data, to ensure the model does not learn from the noise (variability).

Huesken’s dataset has 2431 interactions across 34 mRNA targets. siRNAs from Huesken’s experiments were unmodified and 21nt long, transfected at a fixed 50nM concentration and efficacies were calculated as normalized percentages of reporter inhibition, derived from the reduction of reporter signal in siRNA-treated samples relative to controls and scaled from 0% (least active) to 100% (most active).

The remaining data from non-Huesken experiments (referred to as the *Independent Dataset*) was used to evaluate the model trained on Huesken’s dataset, to test model generalizability.

### Sequence Representation: Z-curves

To train an efficient model it is important to identify appropriate features that represent the data [17]. Previous works on efficacy prediction have identified different ways of representing sequences such as: one-hot encodings, K-mer counting and RNA foundation model embeddings being the most common methods [5], [7], [8], [6], [18].

However, there are several other representations (z-curves, chaos game representations, etc.) that haven’t yet been explored for efficacy prediction, but have been implemented for other use cases such as DNA sequence representations.

We have utilized two such packages that extract various numeric features/representations from RNA sequences-RNAincoder and MathFeatures [19], [17]. RNAincoder extracts several sequence, structure and physico-chemical specific properties whereas MathFeatures use different numeric encodings to represent sequences such as Chaos Game Representations and Complex Networks [20], [21].

We framed efficacy prediction as a classification task, directly distinguishing siRNAs with *High* (efficacy *>* 0.7) versus *Low* efficacy to avoid sensitivity to noisy efficacy measurements and better mirror clinical decision-making. The threshold was chosen for consistency with pharmaceutical pipeline decision making and consistency with previous studies [7, 9]. Class balance was evaluated across multiple threshold values (0.6–0.9), with 0.7 emerging as the optimal threshold for achieving the most balanced class distribution (supplementary section 1). We evaluated our representations using seven classification models: Random Forest, Logistic Regression, XGBoost, Stochastic Gradient Descent, K-Nearest Neighbours, Support Vector Classifier, and Decision Trees. For all models, we retained default scikit-learn 1.7.2 parameter settings as a conservative baseline. The best-performing representation and model combination was then selected for further model explainability.

Supplementary Table 1 describes the different representations we have tested with our model. Upon comparing the various representations and group of features, our model evaluation metrics were highest using Logistic Regression on z-curves [22] (Fig.2). Z-curve was originally introduced as a way to represent DNA sequences in a 3D format. It uses 3 dimensions to identify each nucleotide based on – Weak/Strong hydrogen bond, Purine/Pyrimidine bases, and the presence of Keto/Amino groups in similar positions. Moreover, the original sequence can be reconstructed from the z-curve. Beyond its original role in representing DNA sequences, the Z-curve has also been applied to the representation and analysis of ncRNAs [23], as well as to genome analysis for identifying protein-coding genes [24], among other applications. Our 21nt siRNAs were therefore converted to 1D vectors of length 63 (21×3). Most numeric representations and embeddings cannot reconstruct the original data, which could lead to loss of information in the process of data conversion. One of the important features mentioned by siRNADiscovery was the interaction between RNA and the AGO-2 protein. We have been able to capture this through the representations as well which is demonstrated from our sliding window experiments explained in Results.

### Choice of Cross-Validation

In previous works, 10-fold cross-validation has been the most common method of evaluating the model [5], [8], [6], [7]. However, this method can be biased due to the repetition of a target in both train and test folds. A recent work, siRNADiscovery [9], introduced a new cross-validation method that can prevent data leakage. Their method suggests splitting the data by groups of targets/siRNAs. This means the same target/siRNA is not going to repeat in the train and test folds. Although group based cross-validation makes sure that targets are not repeated in train and test, it can still introduce bias based on the choice of targets in the test set. We have evaluated these biases through a few tests.

K-fold cross-validation assumes, i.i.d (Independent and identically distributed data) to ensure there is no bias in training/testing [25]. We have conducted some tests to identify the possibilty of target (group) based bias in the dataset. We have tested for dependence between ground-truth class labels and target genes using chi-square test, we have tested the dependency between efficacy and target genes using one-way ANOVA and finally we have tested for intraclass correlation (ICC) of the efficacy values across the target genes.

Table 1 demonstrates that when the group order is preserved, the null hypothesis is rejected; however, when the groups are shuffled, it is accepted, implying that the test results depend on the original group structure.

**Table 1.** I.I.D Test to test target-efficacy dependencies confirms that there is significant impact of the group/target order in the dataset.

| Test | Statistics for shuffled group | Statistics for ordered group |
| --- | --- | --- |
| Chi-square P-value | 0.0737 | <0.0001 |
| ANOVA P-value | 0.5149 | <0.0001 |
| ICC | $3.67 \times 10^{-9}$ | 0.1648 |

Next, to test for bias in Group Shuffle Split or the method suggested by Long et al in siRNADiscovery, we have calculated the occurrence of each target in the test set for different train-test split sizes. This is to test the assumption that if a target group has more interaction data as compared to others, it is less likely to be selected in the test set (depending on the test holdout percentage) Figure 1.

**Fig 1.**
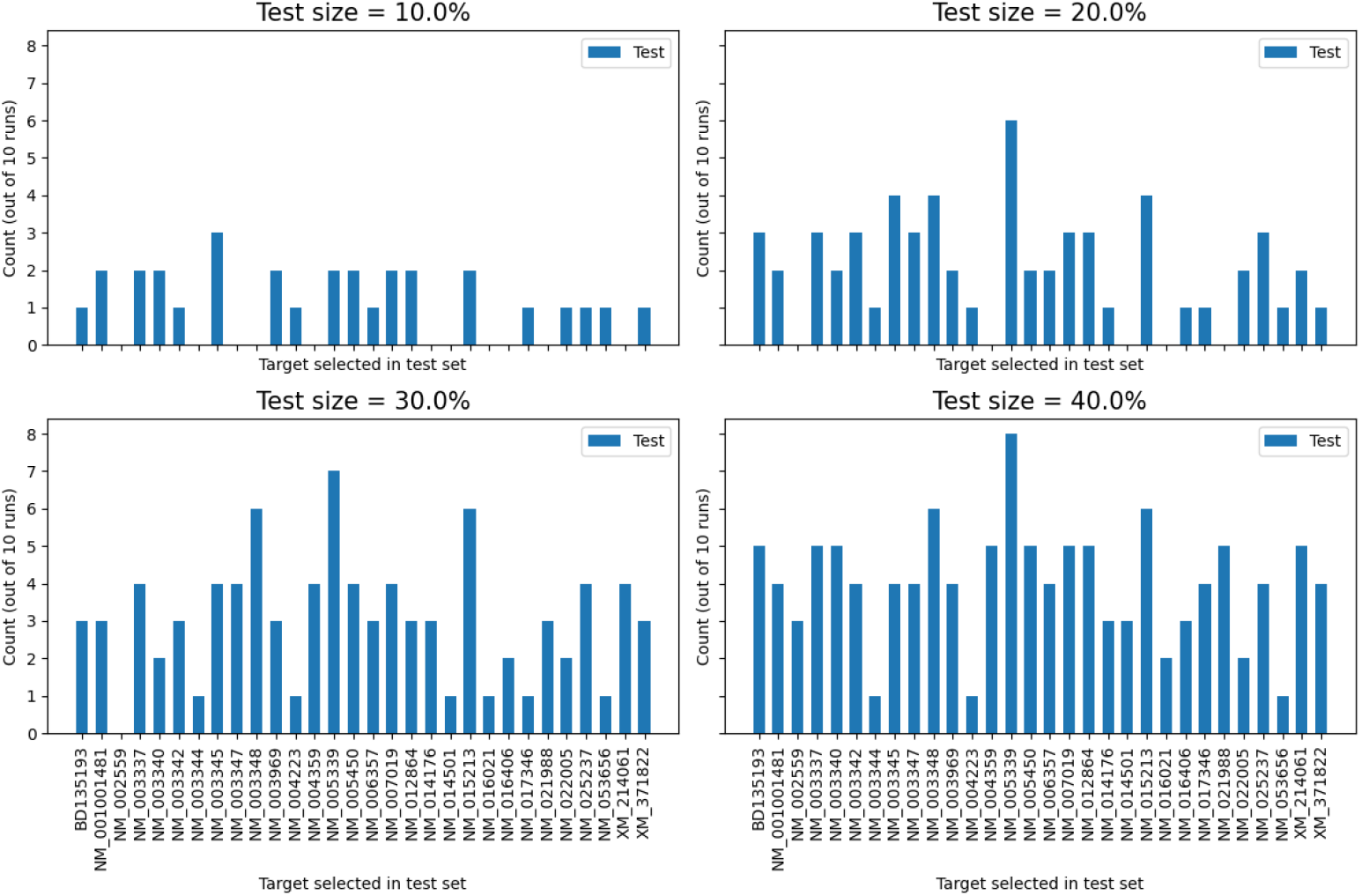
The Group Shuffle Split test target selection is illustrated using various test set percentages, showing that certain targets are excluded from the test set because their data size exceeds the designated test set size.

**Fig 2.**
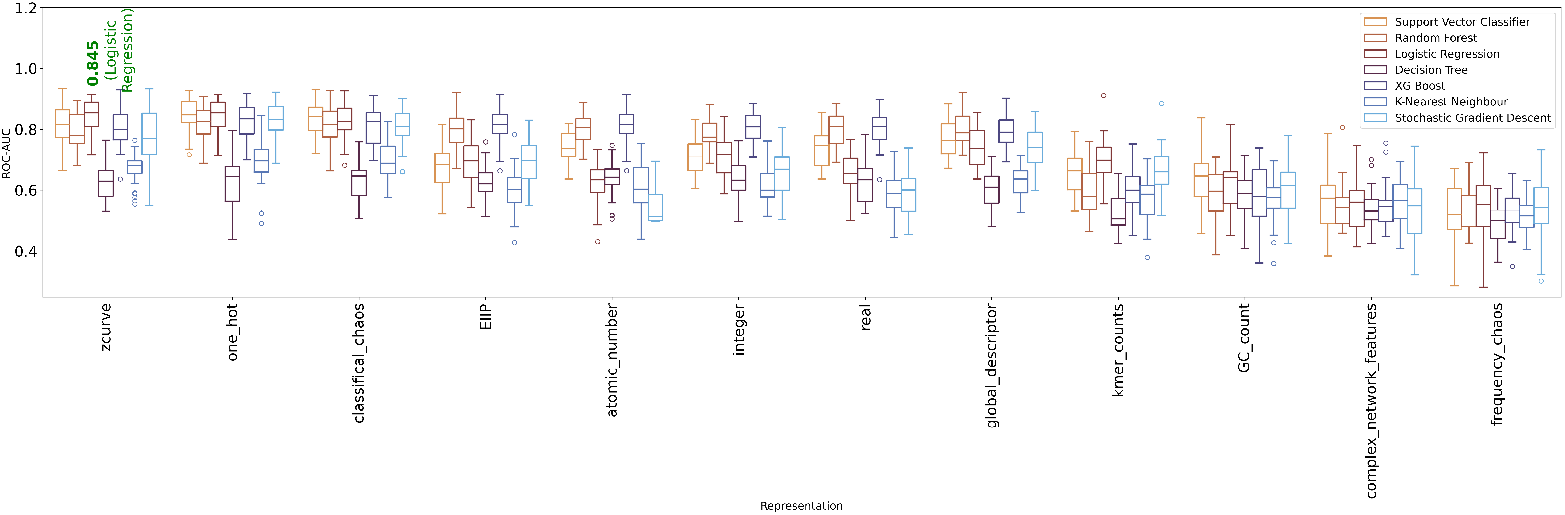
ROC-AUC across models and representations (Supplementary Table 1) shows that the Z-curve achieve the highest performance amongst the methods tested with One-hot encodings and Chaos Game Representations reaching almost to the same values.

Finally, we propose using Leave One Target Out cross-validation, where in each iteration, we shall keep one target gene group out for testing and train on the rest. This method helps us understand the model’s performance on each target and enables us to identify targets that perform lower or do not follow the binding rules typically followed by the rest of the targets of the training group.

### Exploring Important Regions of siRNA

The objective of these experiments was to construct an explainable model capable of capturing the complex mechanisms and key determinants underlying the siRNA knockdown process.

To investigate this, we conducted a sliding-window experiment with varying window sizes, down to a single nucleotide, to determine whether specific regions contribute disproportionately to the knockdown effect.

## Results and Discussion

### Model Cross-Validation Results

Fig.2 presents our model evaluation across several representations and models, showing that z-curve representations achieve the highest performance with Logistic Regression.

Z-curve offers several advantages over one-hot encodings and CGR, such as capturing meaningful chemical features, preserving global positional context, and its low dimensionality makes it well suited for classical machine-learning models. Therefore, we base our subsequent experiments on the z-curve representation. The

Leave-One-Target-Out evaluation yields an average performance of 0.845 (Fig.3), with most targets achieving scores above 0.8. To account for differences in training set sizes, we conducted an additional experiment by fixing the training size to the minimum available across targets. Under this constraint, the model’s performance only decreased to 0.844 (Fig.4), demonstrating its robustness.

**Fig 3.**
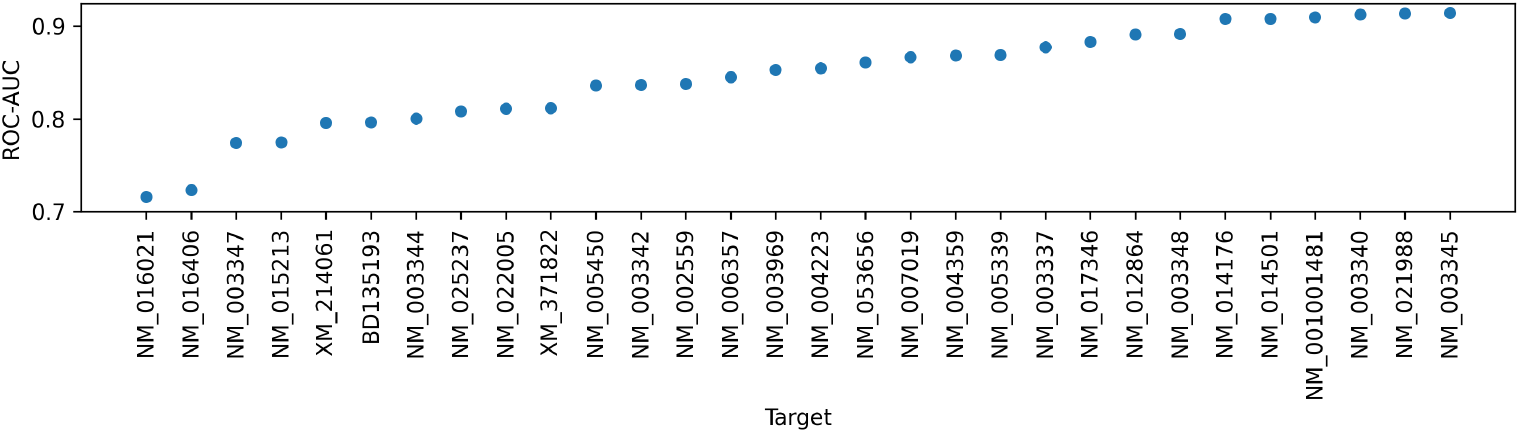
Targetwise results for Logistic regression on Z-curve representations. With an average AUC of 0.845, most targets exhibit performance above 0.80.

**Fig 4.**
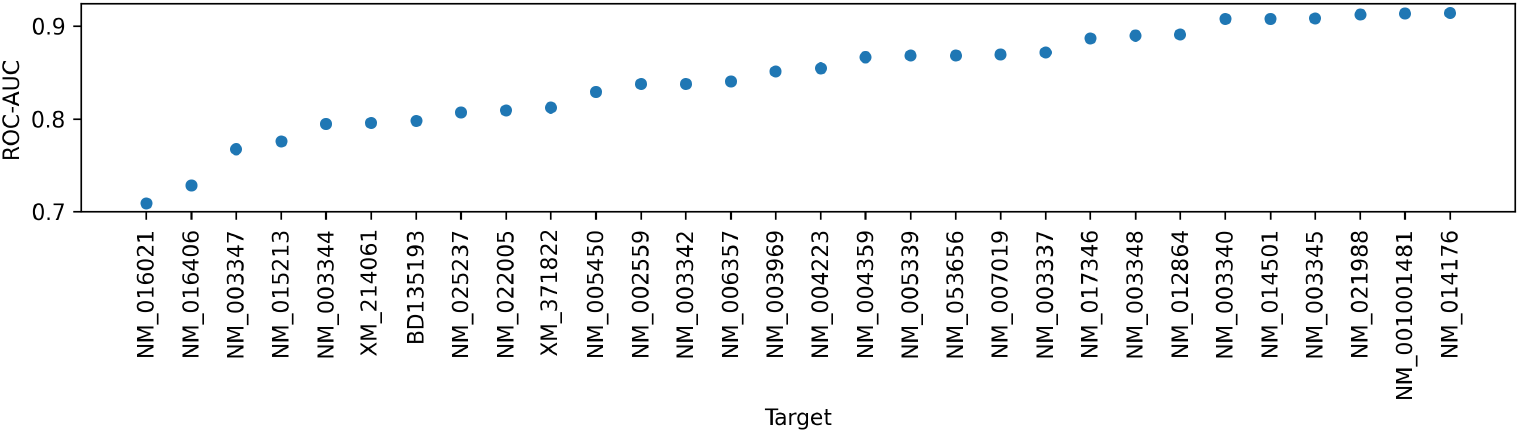
Targetwise results for Logistic regression on Z-curve representations with fixed training size (training size fixed to the smallest value). The reduction in training size had minimal impact on performance, with an average AUC of 0.844.

This cross-validation method is robust, unbiased and provides insights into each target gene’s performance individually unlike K-fold cross validation or Group Shuffle Split.

### Comparison with Other State of the Art Models

As discussed earlier, traditional machine learning models are often dismissed in favor of deep learning approaches due to their typically lower predictive accuracy. However, our results highlight a crucial but sometimes overlooked advantage: computational efficiency. When considering both training and inference time, our model demonstrates a substantial speed advantage, underscoring the importance of learning strong, informative representations rather than relying solely on architectural complexity.

To benchmark against modern deep learning approaches, we compared our method with Oligoformer [7] and siRNADiscovery [9]. These models employ state-of-the-art architectures (transformers and graph neural networks), respectively - and have previously demonstrated superior performance over earlier methods.

For a fair comparison, all models were evaluated using the same

Leave-One-Target-Out cross-validation protocol on Huesken’s dataset and executed on identical hardware. The Leave One Target Out AUCs were then compared with independent 2 sample t-test. As shown in Fig. 5, our model achieves a higher AUC (p-value *<*0.05) than the transformer-based Oligoformer and performs comparably (p-value *>*0.05) to the graph-based siRNADiscovery. Our model’s training and inference time is also ≈ 380 times lesser than siRNADiscovery and ≈ 3070 times lesser than Oligoformer, highlighting that well-designed representations can deliver competitive predictive performance while dramatically reducing both training and inference time.

**Fig 5.**
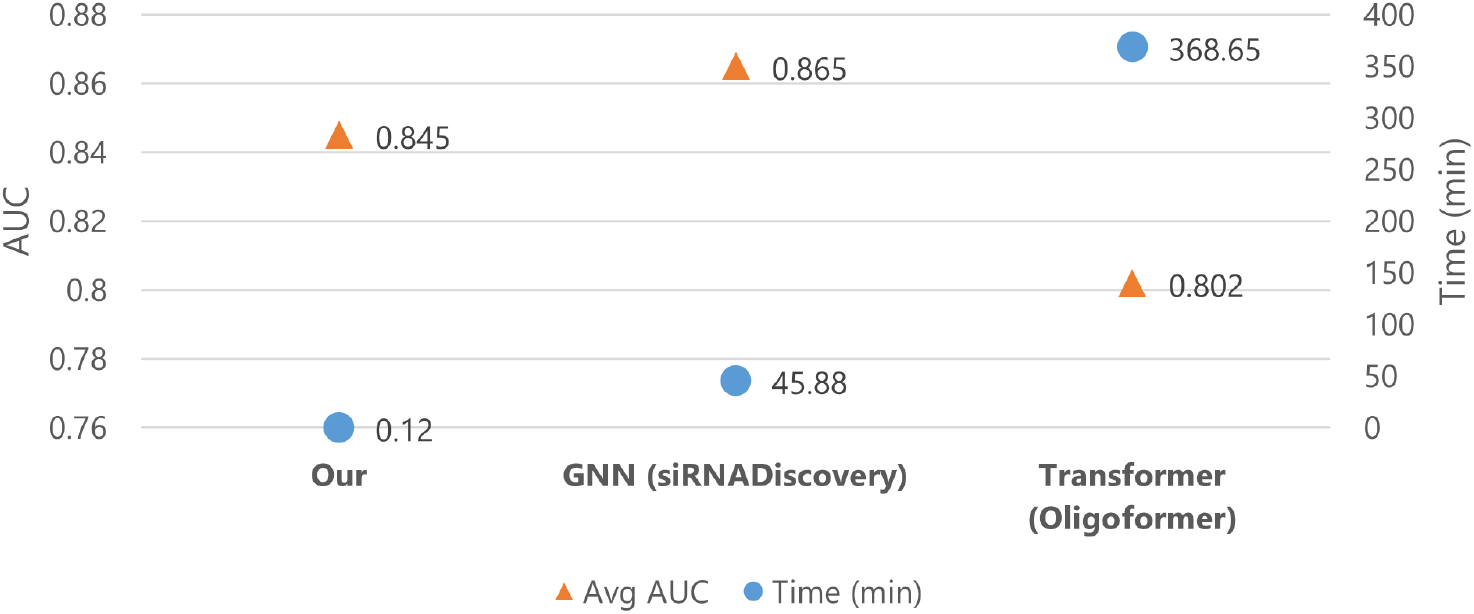
Our model compared with Oligoformer and SiRNADicovery, reveals comparable AUC values with significantly lower compute time (minutes).

This indicates, when combined with the right representations, machine learning models can perform as well as deep models while saving on compute cost and resources.

### Evaluation on Independent Data

We further assessed the robustness of our model by training it exclusively on Huesken’s dataset and evaluating its performance on all remaining datasets. Under this cross-dataset setting, the predictive AUC dropped to 0.753 (Fig. 6), highlighting a substantial loss in generalizability.

**Fig 6.**
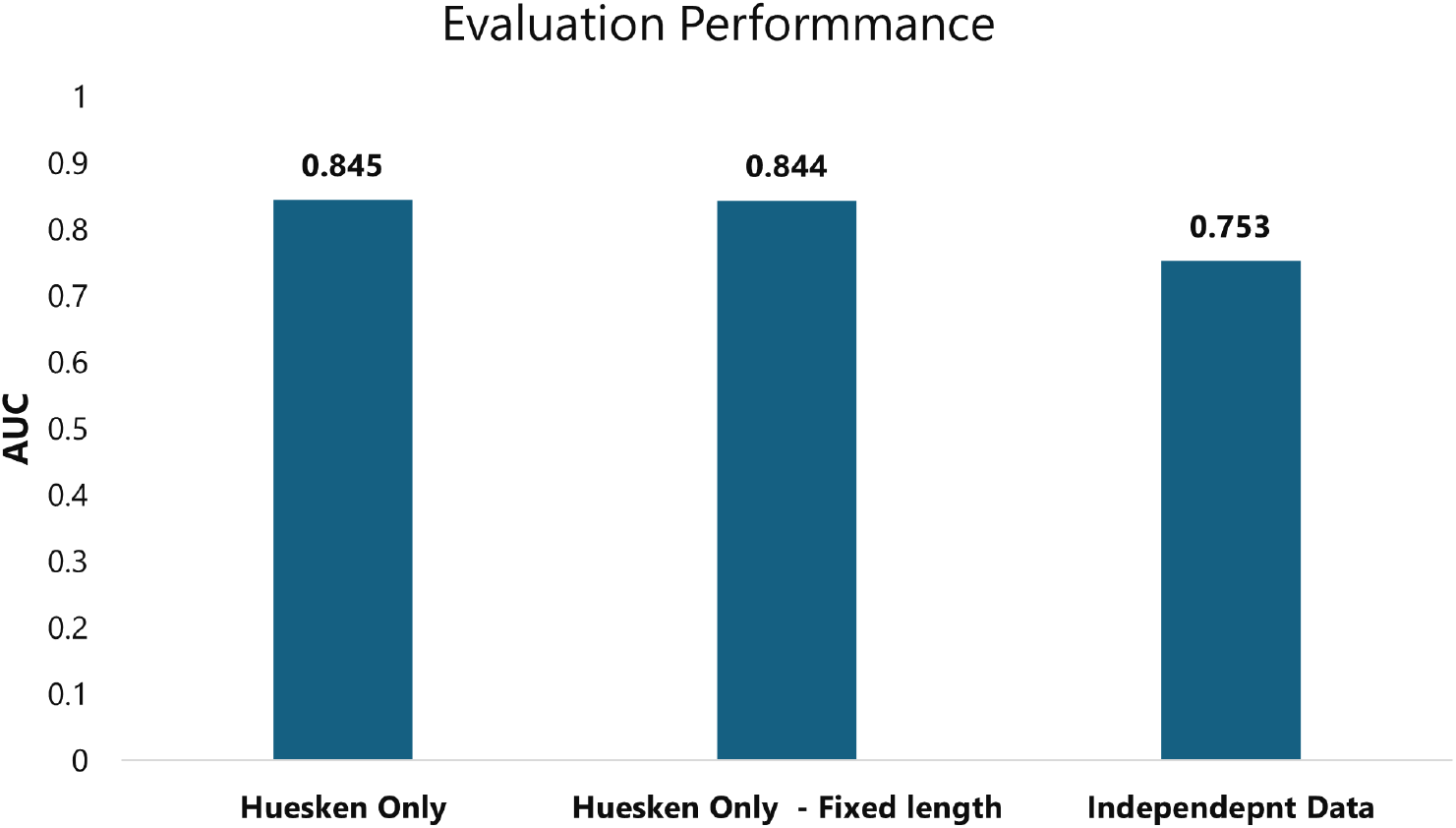
Testing on independent (non-Huesken) data results in a decrease in AUC, suggesting a potential source of noise or error in the evaluation. This drop likely stems from heterogeneity in the experimental conditions across datasets.

This drop likely reflects the model’s limited capacity to generalize beyond the specific experimental conditions it was trained on, as it struggles to account for variability in experimental factors across different datasets measuring the same underlying phenomena. Experimental conditions vary widely, including differences in assay protocols, measurement modalities, siRNA concentrations, and the presence or absence of chemical modifications. For instance, the studies by [12] and [16] incorporate chemically modified siRNAs, whereas others rely on unmodified sequences. Similarly, while most datasets are derived from mammalian cell lines, [15] spans multiple species, introducing additional biological variability. Moreover, knockdown efficiency is quantified using diverse readouts: fluorescence-based assays, protein levels, or mRNA measurements, further compounding inconsistencies.

Collectively, this inter-dataset heterogeneity introduces systematic noise and distributional shifts that impair the model’s ability to generalize beyond its training domain. These findings underscore the importance of explicitly modelling experimental context such as concentration, chemical modifications, and assay-specific factors to better account for variability and improve cross-study robustness. However, the scarcity of experimental data remains a major bottleneck for this aspect.

### Importance of 5^′^ Nucleotide

In our sliding window experiments, we observed that the presence of the first nucleotide within the window led to improved performance even if the window size is one i.e. trained on each individual nucleotide (Fig.7). This was further validated using mRNA sequences, where including the nucleotide that binds to the 5^*′*^ end of the siRNA within the window also resulted in higher performance. These findings confirm that the nucleotide at the 5^*′*^ end of the anti-sense strand is a key contributor to knockdown efficiency, consistent with the results reported by Frank et al. [26]. Their study attributed this importance to the binding of the 5^*′*^ nucleotide to the Ago-2 protein, which facilitates mRNA cleavage. We also observed that some positions such as 2, 7 and 19 were more important than others. The sequence length is important as well, as we observe that adding more information i.e more nucleotides to the training model increases the accuracy however majority of the information comes from the 5 ^*′*^ nucleotide.

**Fig 7.**
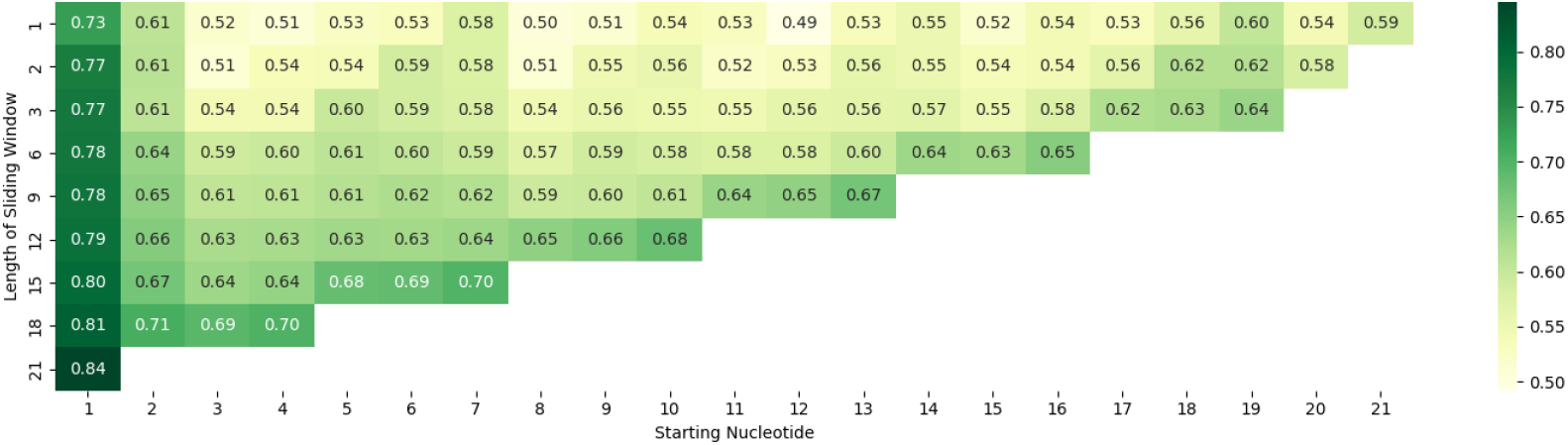
The model is trained with different sliding window lengths across different regions (x-axis shows the starting nucleotide of window). The results highlight the significant contribution of the 5^*′*^ nucleotide. It also highlights a few other positions contributing to a higher performance than the rest. The y-axis indicates the importance of window size i.e. adding more information leads to better performance.

Our model’s predictions using only the 5^*′*^ nucleotide further supported this, showing that A/U at the 5^*′*^ end is associated with higher knockdown, whereas G/C is linked to lower knockdown.

Overall, these experiments revealed several factors influencing the knockdown process, some of which have been demonstrated previously in wet-lab studies and are now corroborated through predictive modelling. Moreover, they highlight the effectiveness of the z-curve representation in capturing both chemical and positional information, enabling the model to learn and extract the relevant underlying binding rules.

### Issues with Data Scarcity and Heterogeneity

Complex deep learning architectures, such as convolutional neural networks or graph-based models, generally require significant training time and computational resources for optimization. In contrast, traditional machine learning algorithms are typically faster, though often at the expense of predictive accuracy. In this work, we evaluated several machine learning models on z-curve representations for the task of classifying high- and low-efficacy sequences. Our best-performing model achieved an

ROC-AUC of 0.845, which is comparable to results reported in the literature, but with considerably reduced computational cost. The ability to train efficiently on new data and generalize to unseen samples highlights the importance of such lightweight yet accurate models. Furthermore, our results demonstrate that suitable sequence representations can enhance the predictive power of even a simple algorithm such as logistic regression.

A well-recognized challenge of deep learning approaches lies in their heavy dependence on large training datasets, which are required to capture complex patterns. Machine learning models, by contrast, can often perform reasonably well with limited data, albeit sometimes at reduced accuracy. To evaluate the data dependency of our approach, we conducted an experiment where the training set size was gradually increased, and the resulting performance was measured on the *Independent Dataset*.

The obtained learning curve suggests that our model is indeed data-bound as seen in Fig.8, and that additional data would likely improve performance further.

**Fig 8.**
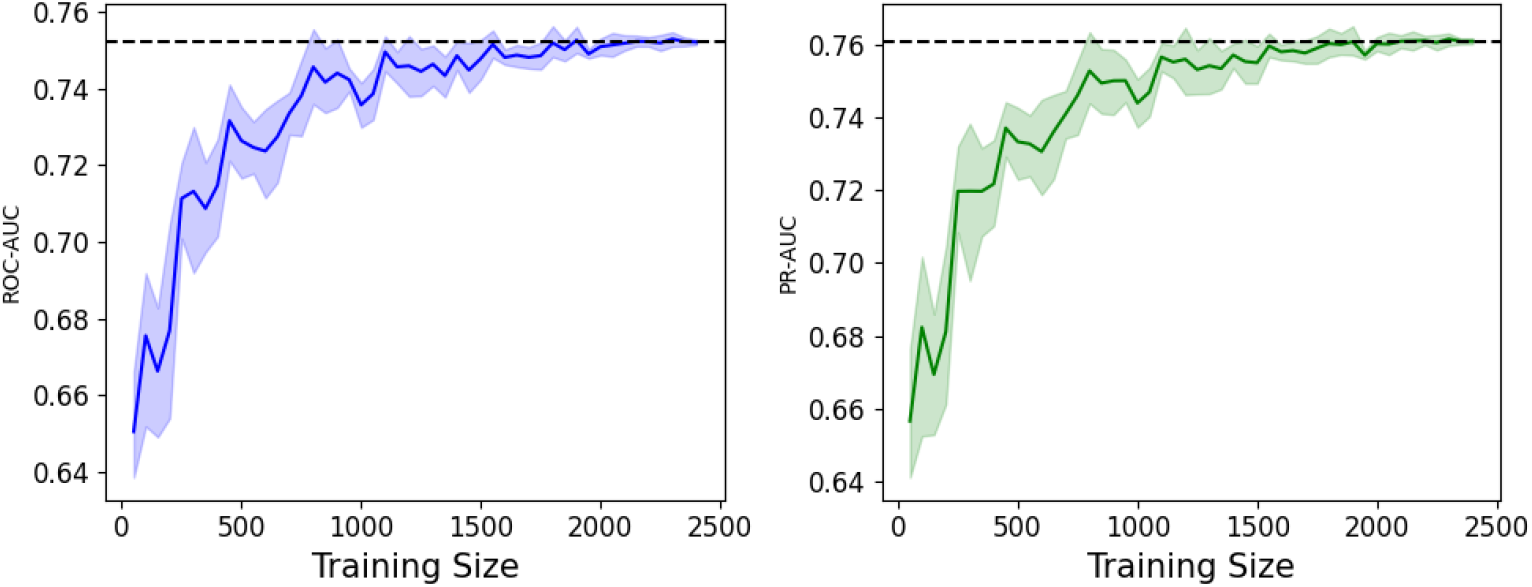
Our model is tested with increasing training size and the results illustrate how increasing data points, increase the AUC values as well suggesting that our model data bound.

However, most publicly available datasets originate from experiments conducted by different research groups, each employing distinct protocols, siRNA designs, and fold-change calculation methods. Merging such heterogeneous data introduces experimental variability, thereby increasing noise and ultimately weakening the model.

## Conclusion

In this work, we propose using Z-curves as an efficient sequence representation method for machine learning tasks. Z-curves transform sequences into three-dimensional numerical representations that capture sequence context and physico-chemical properties, while preserving the original sequence information (i.e., they are reversible). These information rich representations can be used with simple machine learning model to predict siRNA knockdown efficacy.

Our model attains an AUC of up to 0.845 and successfully identifies key siRNA–mRNA binding determinants, including the critical role of the 5^*′*^ nucleotide in mediating the knockdown process. This demonstrates that our model is both accurate and interpretable.

We aim to benchmark the cross-validation approaches applied in siRNA experiments, demonstrating the bias introduced by methods like k-fold cross-validation or group shuffle split. As a more stringent alternative, we propose using Leave One Target Out cross-validation, which yields clearer insights into individual targets.

Finally, while our current dataset is insufficient to fully address these challenges, we demonstrate that integrating heterogeneous experimental datasets is problematic due to potential noise introduced by variability in experimental protocols. Furthermore, we underscore the critical need for larger and more consistent datasets to enable the development of reliable and accurate models, which remains a key limitation in the present study.

## Key Points

1. We propose Z-curve as an efficient representation of RNA sequences, which can be used as input features to the prediction model.
2. We have addressed the data leakage issue with previously implemented cross-validation methods by using target-based data splitting. We propose using Leave One Target Out Cross Validation.
3. Finally, we use this model to explain the important regions of siRNA responsible for effective knockdown.

## Acknowledgments

We gratefully acknowledge the Engineering and Physical Sciences Research Council (EPSRC), the Rosalind Franklin Institute, and the School of Chemistry at the University of Edinburgh for their financial support of this study.

