## Supplemental Data for "Benchmarking siRNA Prediction: The Role of Representation and Validation Strategies"

### Supplementary Data

#### 1 Huesken Data Distribution

Figure 1 shows how label distributions vary across different threshold choices. Although a 70% (0.7) threshold is commonly used as the standard in the literature, we also examined distributions across multiple thresholds, which clearly indicate that 0.7 provides the best balance.

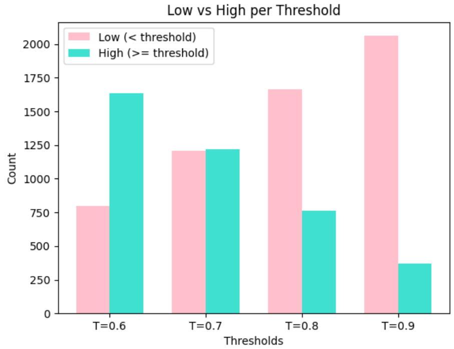


**Fig 1.** Distribution of Classes/Labels across different choices of thresholds on Huesken's dataset.

#### 2 Statistical Test of Model Comparisons

We implemented Leave One Target Out Test on the Huesken's dataset for our model in comparison with siRNADiscovery(Graph neural networks) and Oligoformer(Transformer based) models. Results demonstrate equivalent performance with siRNADiscovery with no significant difference and signifcantly higher performance that Oligoformer.

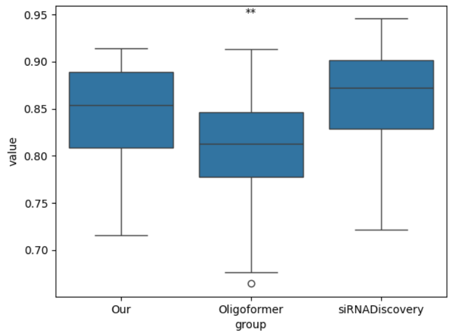


**Fig 2.** ROC-AUC for Leave One Target Out experiment across the three models. Results show how Our model significantly outperforms Oligoformer while performs almost as good as siRNADiscovery without any significant difference in perfromance.

#### 3 Choice of Cross-Validation

We have evaluated the Huesken's dataset with k-Fold(k=10) cross-validation which is commonly used in the literature and Leave One Target Out with fixed training size.

While 10-fold yields stable but lower median performance, leave One Target Out reveals higher variability alongside a higher median, suggesting that the model generalizes well to many groups but fails on a subset. These results highlight inconsistent generalization across groups and demonstrate that standard cross-validation may mask group-specific performance differences, whereas group-aware evaluation provides a more realistic assessment.

To further assess whether the variability in group performance under the Leave-One-Target-Out approach is driven by target-specific properties or underlying data distributions, we examined the Pearson correlation between ROC-AUC and target data size ($-0.012$), as well as between ROC-AUC and the target class ratio (percentage of class 1/high knockdown; 0.060). Both correlations are close to zero, suggesting that the observed variability is more likely due to differences in target behavior rather than data size or class balance. These results further emphasize on the importance of using group-aware splitting.

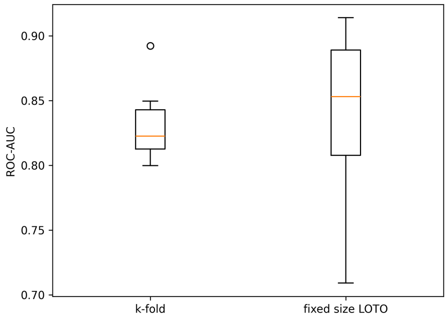


**Fig 3.** The broader distribution observed with LOTO, versus the stable AUC range from 10-fold cross-validation, indicates that 10-fold under-represents group-level heterogeneity.

### 4 Representations

**Table 1.** Methods used to represent sequences

| **Representation** | **Source/Package** | **Description** |
|---|---|---|
| GC-count | RNAincoder | Distribution of G/C across the sequence |
| Global Descriptor | RNAincoder | Nucleotide composition, transition and distribution representation of a sequence |
| Frequency Chaos | MathFeatures | Sequences to fractal-like images based on k-mer frequencies |
| Complex network | MathFeatures | Betweenness, assortativity, average degree, average path length, minimum degree, maximum degree, number of edges, degree SD, frequency of motifs, clustering coefficient |
| Classifical Chaos | MathFeatures | Jeffery's Classical Chaos Game Representation |
| Integer | MathFeatures | $X(i) = \begin{cases} 3, & \text{if } X(i) = G \\ 2, & \text{if } X(i) = A \\ 1, & \text{if } X(i) = C \\ 0, & \text{if } X(i) = T \end{cases}$ |
| Z-curve | MathFeatures | $X_1(i) = \begin{cases} X(i-1)+1, & \text{if } X(i) = A \vee G \\ X(i-1)-1, & \text{otherwise} \end{cases}$ $X_2(i) = \begin{cases} X(i-1)+1, & \text{if } X(i) = A \vee C \\ X(i-1)-1, & \text{otherwise} \end{cases}$ $X_3(i) = \begin{cases} X(i-1)+1, & \text{if } X(i) = A \vee T \\ X(i-1)-1, & \text{otherwise} \end{cases}$ |
| One-hot | Custom | $X_1(i) = \begin{cases} 1, & \text{if } X(i) = A \\ 0, & \text{otherwise} \end{cases}$ $X_2(i) = \begin{cases} 1, & \text{if } X(i) = T \\ 0, & \text{otherwise} \end{cases}$ $X_3(i) = \begin{cases} 1, & \text{if } X(i) = G \\ 0, & \text{otherwise} \end{cases}$ $X_4(i) = \begin{cases} 1, & \text{if } X(i) = C \\ 0, & \text{otherwise} \end{cases}$ |
| Real | MathFeatures | $X(i) = \begin{cases} -0.5, & \text{if } X(i) = G \\ -1.5, & \text{if } X(i) = A \\ 0.5, & \text{if } X(i) = C \\ 1.5, & \text{if } X(i) = T \end{cases}$ |
| EIIP | MathFeatures | $X(i) = \begin{cases} 0.0806, & \text{if } X(i) = G \\ 0.1260, & \text{if } X(i) = A \\ 0.1340, & \text{if } X(i) = C \\ 0.1335, & \text{if } X(i) = T \end{cases}$ |
| K-mer counts | Custom | Code RNA sequences through the occurrence frequencies of k neighboring nucleic acids |
| Atomic Number | MathFeatures | $X(i) = \begin{cases} 78, & \text{if } X(i) = G \\ 70, & \text{if } X(i) = A \\ 58, & \text{if } X(i) = C \\ 66, & \text{if } X(i) = T \end{cases}$ |
